# Haplotag: software for haplotype-based genotyping-by-sequencing analysis

**DOI:** 10.1101/031013

**Authors:** Nicholas A Tinker, Wubishet A Bekele, Jiro Hattori

## Abstract

Genotyping-by-sequencing (GBS) and related methods are based on high-throughput short-read sequencing of genomic complexity reductions followed by discovery of SNPs within sequence tags. This provides a powerful and economical approach to whole-genome genotyping, facilitating applications in genomics, diversity analysis, and molecular breeding. However, due to the complexity of analysing large data sets, applications of GBS may require substantial time, expertise and computational resources. Haplotag, the novel GBS software described here, is freely available and operates with minimal user-investment on widely-available computer platforms. Haplotag is unique in fulfilling the following set of criteria: (1) operates without a reference genome; (2) can be used in a polyploid species; (3) provides a discovery mode and a production mode; (4) discovers polymorphisms based on a model of tag-level haplotypes within sequenced tags; (5) reports SNPs as well as haplotype-based genotypes; (6) provides an intuitive visual “passport” for each inferred locus. Haplotag is optimized for use in a self-pollinating plant species.

---

**Summary (100 words):** This report describes and makes freely available a novel software application designed to analyze and report results of genotyping-by-sequencing. The software takes a novel approach to discovery and validation of loci based on tag-level haplotypes within clusters of aligned tags that may contain multiple paralogous loci. Output from these analyses are reported in multiple formats, including an intuitive passport showing discovered loci and genotypes within each cluster.

Genotyping-by-sequencing (GBS: Elshire et al. 2011) and similar methods (e.g. RAD: Miller et al. 2007) have become important strategies for whole genome genetic diversity analysis and related studies in many plant and animal species. The objective of these strategies is to resequence a representative fraction of the genome of many individuals, and thereby determine the genotypes of those individuals at loci where sequence variants exist. Methods are based on high-throughput short-read sequencing of enzymatically-constructed genomic complexity reductions, followed by discovery of SNPs within sequence tags. While GBS is powerful and economical, it is also complex: requiring the barcoding and multiplexing of samples, the deconvolution of large data files, the alignment of short reads (tags), and the discovery and filtering of SNPs. The application of GBS in large and complex genomes is especially challenging because of the confounding presence of multiple paralogous loci (especially in polyploids), and often, the absence of a complete reference genome.

There are several available bioinformatics pipelines for GBS analysis, including Stacks (Catchen et al. 2011), TASSEL (Glaubitz et al. 2014), UNEAK (Lu et al. 2013) and other custom-designed pipelines (e.g. Sonah et al. 2013; Poland et al. 2012). Most pipelines require or benefit from a reference genome, while UNEAK is designed specifically to operate independently from a reference genome and Stacks has the ability to run with or without a reference genome. Stacks is a flexible and integrative set of tools tool that produce many types of output and can be customized for many genetic scenarios. Stacks also provides a unique web-based interface for inspection of results and quality control: a feature that is useful in tuning the many parameters of GBS analysis such that they produce results that are appropriate to the genome and the genetic population. However, Stacks requires a Unix-like computer environment and a significant investment of effort in building and maintaining a pipeline, and the web-based interface requires a relational database and web server. Most other GBS pipelines also require the installation of third party programs (e.g. to align sequences) while UNEAK requires only the installation of a JAVA run-time environment.

To our knowledge, UNEAK and the customized scripts described by Poland et al. (2012) are the only existing pipelines that will handle data from polyploids in the absence of a reference genome. Both pipelines achieve this by using a population filter to rejects SNPs that fail to segregate with the expected genetic ratio in the population under analysis. Because UNEAK can be run on any computer platform with adequate resources, it has been popular among researchers studying species where no reference genome is available. However, the UNEAK pipeline excludes all SNPs that belong to multi-locus series, SNPs from tags containing multiple SNPs, or SNPs with more than 2 alleles. In our experience with GBS in hexaploid oat (Huang et al. 2014) UNEAK excluded at least 30% of potentially useful SNPs that were discovered by an alternate customized pipeline. Furthermore, the developers of UNEAK (personal communication) have indicated that no further development of UNEAK will be performed.

With high-density genotyping comes the possibility to analyse data based on haplotypes and the ability to impute missing data (Swarts et al. 2014) which may be of particular importance in GBS analyses where incomplete data are prevalent. Genome wide association studies (GWAS) based on haplotypes could also allow the discovery of cryptic QTL associations that have eluded analysis based on single SNPs (Lorenz et al. 2010). Because GBS data are acquired from sequenced fragments that often contain multiple SNPs, direct information about localized ‘tag-level’ haplotypes are available within a GBS pipeline. However, to our knowledge, no GBS pipeline is able to examine the segregation of haplotypes in the application of a population filter, nor does any software provide a simple method to access or examine haplotypes directly in an output file. Since accurate haplotype inference normally requires a reference genome, the availability to extract haplotypes directly from within GBS fragments could be of particular interest in a species where no reference is available.

Our objective was to develop user-friendly GBS software that operates with minimal user-investment on widely-available computer platforms. Additionally, we intended this software to meet the following requirements: (1) to operate without the requirement for a reference genome; (2) to operate in a polyploid or duplicated genome, distinguishing paralogous loci when an appropriate population filter is available; (3) to provide a discovery mode as well as an efficient production mode for scoring previously-discovered loci; (4) to discover polymorphisms based on models of segregating tag-level haplotypes within GBS sequenced tags; (5) to report results in a variety of formats, including SNP- and haplotype-based genotypes, and (6) to provide an intuitive “passport” for each inferred locus, enabling visual inspection and validation of discovered GBS loci.

## Materials and Methods

Software named ‘Haplotag’ was written in the Pascal programming language, implemented as Free Pascal (freepascal.org) within the Lazarus programing environment (lazarus-ide.org). Both of these programming packages are open source, available on multiple platforms, and actively supported by developer communities. Most algorithms within Haplotag were written to operate in parallel when executed on a computer with multiple processors. The code was compiled for the Windows 64-bit environment (Microsoft, Redmond WA) and tested with Windows XP, 7, 8, and 10 and server 2008. Haplotag was tested on many different computers, but evaluations reported below were executed on a computer running Windows server 2008 with two Intel (Santa Clara, CA) Xeon X5670 processors running at 2.93 GHz. Each processor had six cores, and each core was divided into 12 threads (total 24 threads). The test machine contained 96 GB RAM, but all reported analyses were confirmed to run within 24 GB RAM. All input and output data resided on a locally-attached 500GB disk, since prior experience indicated reduced performance when reading and writing to a network drive. Small projects, as well as the demonstration files described below, will run on most ordinary desktop computers, but will require a 64-bit operating system.

Haplotag was evaluated using a set of small simulated demonstration files as well as on the full set of primary GBS reads from oat described by Huang et al. (2014). The later data contained 894 taxa consisting of 360 diverse oat lines and 534 mapping progeny from six bi-parental populations. Both Haplotag and the UNEAK pipeline were run with a minimum merged tag count of 50, which is higher than the threshold used in the earlier work due to subsequent optimization. Output from both pipelines was filtered across the full population to maintain markers for which genotype calls were ≥ 50% or ≥ 80% complete, heterozygosity was ≤ 10%, and minor allele frequency was ≥ 5%. The error detection threshold in UNEAK was set to 0.02. Additional filters for Haplotag included a maximum base difference of 3 for aligning tags, a maximum of 9 tags per cluster, a maximum heterozygote frequency on a haplotype basis of 0.25, and a maximum tolerance for tri-zygotes and multi-zygotes of 1% and 0%, respectively.

### Terminology

When referring to SNPs, we use of the terms ‘SNP locus’ (a specific base pair) and ‘SNP alleles’ (the variant bases found at a SNP locus). We then define a ‘tag-level haplotype’ as the combined set of SNP alleles that must exist on a single chromosome due to their recovery in the sequence of a single GBS tag. Although the term haplotype implies the existence of multiple loci, we essentially treat haplotypes as multiple alleles at a single composite locus, which we refer to as a ‘Haplotag locus’, and inferences are made under the assumption that the recombination rate within a tag is negligible. The term ‘heterozygosity’ is used when applying a filter that rejects an inference that two or more haplotypes exist at the same Haplotag locus if those haplotypes occur together more frequently than they would be expected to based on the assumed heterozygosity in the population.

### Data and software availability

Data analysed in this report were deposited in the NCBI short read archive (http://www.ncbi.nlm.nih.gov/sra/) under project accession number SRP037730, and the GBS key for analysis was available in Table S4 of Huang et al. (2014). Supplemental files include: the Haplotag manual (S1), and sample output (S2 and S3). Haplotag is available as an executable distribution for recent versions of Windows 64-bit environments (XP, and versions 7 through 10). The distribution can be obtained from the site http://haplotag.aowc.ca/ which provides a download links for a compressed file that contains the Windows executable, a user manual (also in S1) and demonstration files. Future updates will be maintained at this site, and a voluntary registration is provided to monitor interest in this software and to enable announcements regarding major revisions. The Pascal source code was made available to reviewers of this work, and will be provided by request on an as-is basis for any noncommercial use based on an open source license. The source code is expected to be compatible with any operating system where a Free Pascal compiler is available, although minor modifications to the code may be required to adapt it for the file systems of other operating environments.

## Results and Discussion

### Software execution

The operation and function of Haplotag is described in the accompanying manual (S1) which references a set of small simulated input files for demonstration purposes. The input files are archived within the software distribution archive. When extracted, the demonstration files fall within three separate subdirectories, each containing a complete self-contained set of demonstration files for one of three primary modes in which Haplotag can operate. Within each subdirectory is a master input file with the default name *“HTinput.txt”* which contains all relevant parameter specifications as well as a set of pipeline commands that Haplotag will follow in the order listed. Based on these commands, Haplotag can read and process data from three starting points (figure 1) representing the three modes of operation.

**Figure 1.**
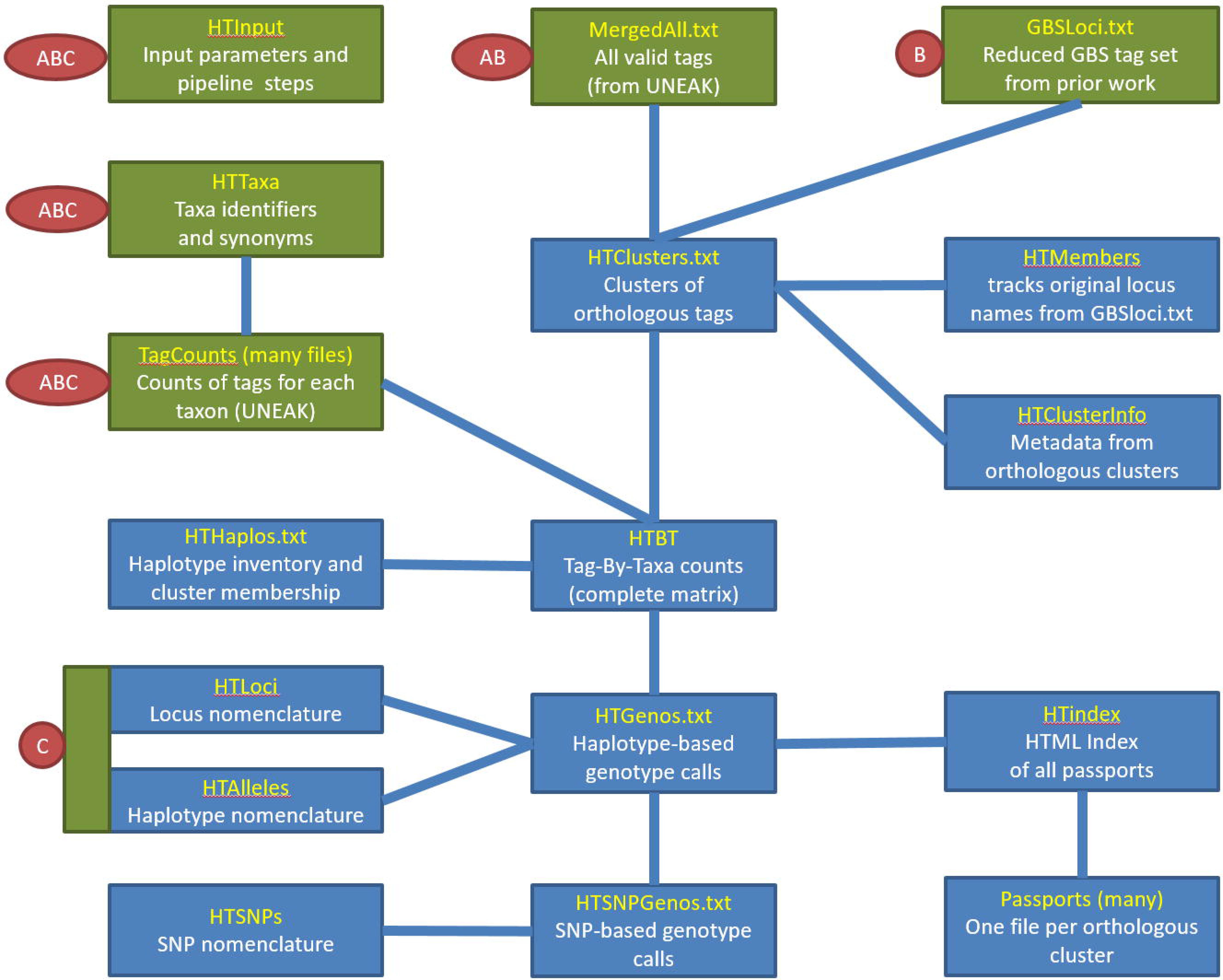
Flow chart showing input files (green), output files (blue) and dependencies (connecting lines) associated with ‘Haplotag’ GBS discovery software. Default file names are shown in yellow, and are normally appended by “.txt” in the Windows file system. Three alternative pipelines (A, B, and C) are available, with required input labeled for each. The cluster discovery pipeline (A) and the haplotype discovery pipeline (B) start by aligning a complete inventory of tags (A) or a reduced inventory of tags from prior work (B) to produce clusters. In (B), the complete inventory is then re-aligned against this template to increase the sampling of new haplotypes. A complete tag-by-taxa matrix of tag counts (HTBT) is then formed for all tags belonging to clusters of two or more tags. Other output files are then created based on haplotype model fitting. In the production pipeline, only the files labelled by (C) are required, since genotyping is based on counting copies of haplotype-tags in the output files from previous discovery work.

There is currently a requirement to run part of the UNEAK GBS pipeline prior to running Haplotag in order to de-convolute the raw barcoded sequence data, produce a tag count file for each sample, and write a merged tag count file for the entire project. The UNEAK pipeline executes these steps very efficiently, thus the replacement of this functionality was not a priority. The current Haplotag distribution provides a small helper utility to assist users in writing the UNEAK script and converting binary output to the text files required by Haplotag. A standalone replacement for UNEAK is being developed which may allow the analysis of tags longer than 64bp, but this tag length is a current limitation of both UNEAK and the current version of Haplotag. Sequencing data with short reads of 100bp is ideal for this type of analysis, since the barcode may occupy op to the first 10 bases, and this allows truncation of lower quality bases at the 3’ end of the read. Reads of longer than 100bp can be analysed, but the tags will be truncated at 64 bases.

The cluster discovery mode (Figure 1A) is designed for applications where complete de-novo SNP discovery is required. This de-novo clustering step is multi-threaded, but it may still run slowly on very large data sets. The haplotype discovery mode (Figure 1B) reduces the scale of analysis by seeding the clusters with a set of pre-determined tags. This feature is useful for maintaining the legacy nomenclature of reference sequences from prior GBS analyses. It could also be used to seed the alignment of clusters using predicted fragments from a sequenced genome. Alternatively, this step could incorporate consensus sequences from an alternate or more efficient clustering algorithm. The production mode (Figure 1C) is designed for applications where SNPs and Haplotypes have already been discovered by Haplotag using a large, diverse and representative population, and where the objective is to genotype new samples while maintaining exactly the same nomenclature of loci, haplotypes, and SNPs. No new haplotypes will be discovered in production mode, so it is not recommended for an application where the diversity of new taxa falls outside of the diversity where the model was built.

What distinguishes Haplotag from other GBS pipelines is the treatment of the tags as haplotypes, and the development of locus models using a population filter to validate the diploid segregation these haplotypes. Prior to model discovery, tags are deliberately over-aligned into clusters that potentially represent multiple paralogous loci. Then Haplotag tests every possible combination of haplotypes within each cluster to identify mutually exclusive groups of haplotypes that behave as single Haplotag loci. This model testing is based on a population filter, which specifies threshold parameters for maximum heterozygosity, minimum and maximum allele frequency, and genotype-completeness (minimum proportion of non-missing genotypes). The result can be a single Haplotag locus within a single cluster, or multiple Haplotag loci within the same cluster. The latter is common in polyploid or recently duplicated genomes. Results of locus prediction and genotype scoring are summarized within a single passport file for each cluster (see below). Although the model selection within clusters does not incorporate sequence divergence, the population filter invariably identifies Haplotag loci in which haplotypes diverge less within the locus than they do among other loci within the same cluster.

### Software function, as illustrated by passport files

Another important and unique feature of Haplotag is the automated production of a ‘passport’ file for each cluster. This is illustrated by one passport from the analysis of the included demonstration data (Figure 2). Passport files are formatted in plain HTML, such that they can be viewed in any web browser. They are indexed in a master HTML file which can also be opened and searched in any browser. While these files can be opened directly from a local disk, they could also be uploaded to a website in order to provide external access to the results of an analysis. Individual passport files can be inspected to determine if program parameters are appropriate, or to explore the metadata and genotypes of specific Haplotag loci. In our experience, these files also serve as intuitive graphical presentations that can assist in explaining the GBS concept and the program function to a lay audience.

**Figure 2.**
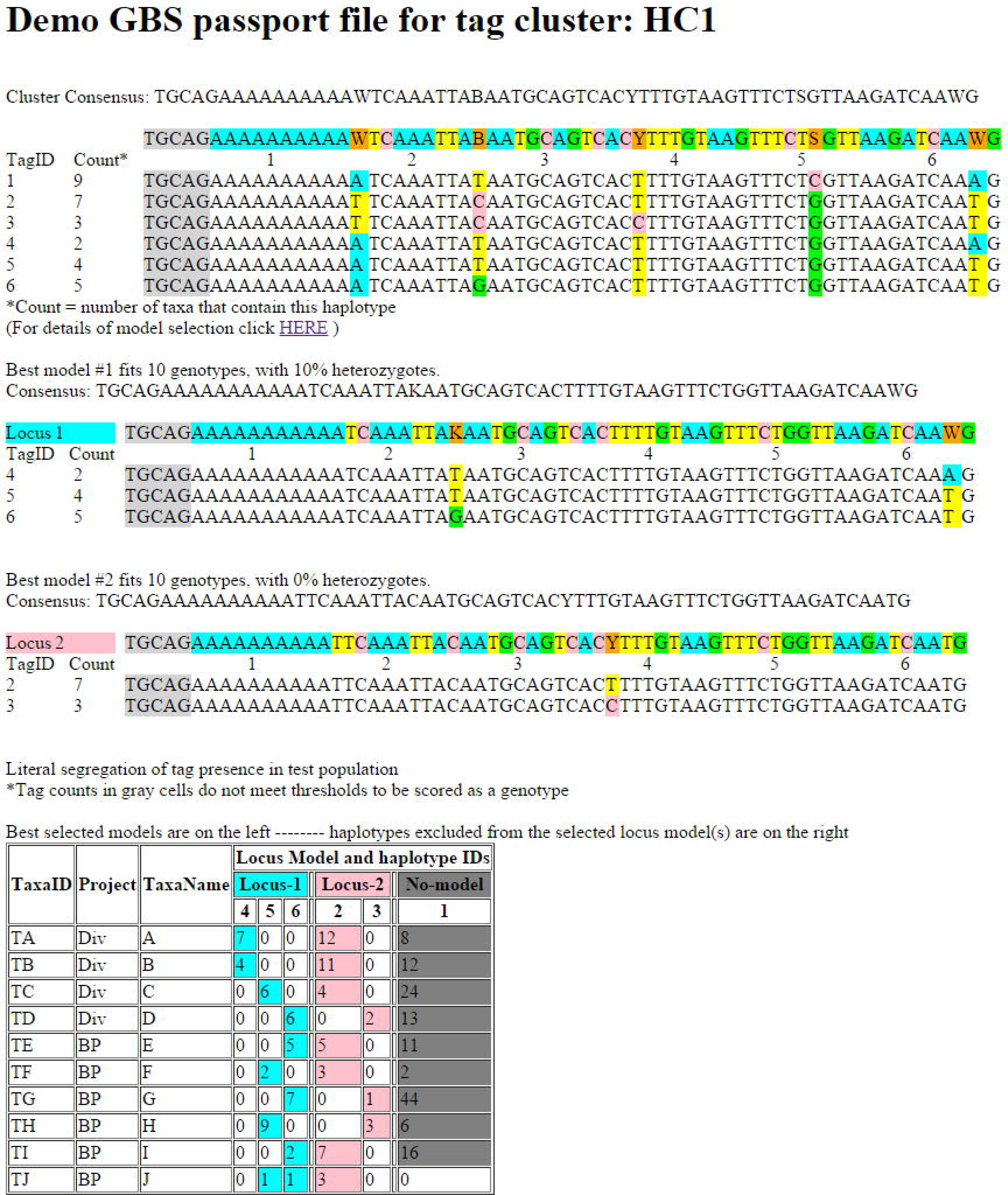
Passport file produced by Haplotag from simulated demonstration files. Here, six tags (potential haplotypes) are identified at the top. After model fitting by population-based filtering, two locus-models are selected. When Haplotag is run in ‘verbose’ mode, the details of model selection are written in a separate file (see S2). Locus-1 contains three haplotypes and Locus-2 contains two. SNP positions are identified by color. The table at the bottom of the passport shows the tag counts at the presumed haplotypes within each locus. Counts greater than one are shaded, indicating that they are scored as “present”.

For example, in Figure 2, we would first explain that the six sequences at the top (TagID 1 to 6) constitute all of the unique 64-base tags from the experiment that formed a single cluster. Potential SNPs in this cluster are highlighted, and counts of each tag are shown at the left. We would then explain that the species from which these tags are generated is polyploid, such that we suspect these tags may come from more than one locus. We might then click on the “details of model” link (which would open table S2) to illustrate how Haplotag has inspected all 57 possible combinations (“models”) of two or more tags from the available six tags. This step is referred to as a “population filter”, since it allows the exclusion of inappropriate models based on whether the tags in a model segregate in a diploid manner within the tested population. Parameters for population filtering, reported at the bottom of the details page (S2) include genotype-completeness, allele frequency, and heterozygosity. Here each model was evaluated based on whether it would pass this filter (yes or no). Next, the acceptable model having complete data for the greatest number of taxa (Model 42 in S2) was assigned as ‘Locus-1’. All models that overlapped with Model 42 were then removed, and remaining acceptable models were inspected. Of these, the next best model was assigned to a Haplotag locus (in this case, Model 48 is assigned as ‘Locus-2’). The above process is iterated indefinitely until no acceptable models remain. We would then point out that ‘Haplotag Locus-2’ contains only one SNP Locus, and thus, two haplotypes while ‘Locus-1’ contains two SNPs, which could theoretically form four haplotypes, of which three haplotypes were observed. In practice, it is very rare to observe four haplotypes at a single Haplotag locus with two SNP loci, as this would imply two mutation events at the same SNP locus, or a rare recombination event between two SNP loci in the same tag.

We would then draw attention to the inferred genotypes and segregation of these five combined haplotypes at two putative Haplotag loci within the population of taxa, which are shown in the table at the bottom of the passport (Figure 2). In this idealized example, the genotypes of all 10 taxa are complete at both accepted Haplotag loci. The numbers in each cell show the total counts of tags observed for each taxon under each haplotype within a selected Haplotag locus. Those with non-zero counts for two (or more) haplotypes (e.g. Taxa TJ, under Locus 1) are scored as heterozygotes. These inferred genotypes are written to a simple text-based file called “HTgenos.txt”. Since many programs for genetic analysis cannot read haplotypes, an alternate genotype file is written where genotypes are defined by SNP locus calls from within the Haplotag loci. In the example in Figure 2, three SNP locus calls would be written, with ‘Locus-1’ being converted to two SNP loci, identified by their SNP positions within the Haplotag loci. Nomenclature output files are also written, such that all dependencies are represented in a hierarchical naming system. These files are designed with shared fields such that they could easily be loaded into a relational database designed for this purpose.

### Parameter selection

It is well known that results of SNP identification, especially in a polyploid without a reference genome, are highly dependent on methods and parameters (Huang et al. 2014). As with other methods for SNP identification, there is no formal way to optimize the selection of model parameters within Haplotag. However, parameters need to be selected carefully, possibly using iterative testing, in order to obtain good results and avoid artefacts. In our experience, the best results from Haplotag are obtained when it is run across a large composite base population consisting of a mixture of bi-parental populations and diverse taxa representative of target germplasm. The bi-parental populations will allow validation of Mendelian segregation and mapping of the polymorphisms, while the diversity samples will ensure discovery of alternate haplotypes. The parameters used for the oat data presented below were based on recursive optimization for this type of experiment. If bi-parental populations are analyzed, then the minimum allele frequency filter can be raised appropriately. If the analysis is restricted to a single bi-parental population, then the filter could be set to achieve a specific chi-square cutoff. Setting the maximum heterozygote frequency to a low value is very useful to exclude non-Mendelian models, but this can only be applied effectively within inbred lines where the expected heterozygote frequency is significantly lower than 50%.

### Evaluation of Haplotag using data from hexaploid oat

Data from 894 taxa reported by Huang et al. (2014) were reanalyzed to compare performance and output of Haplotag to that of the UNEAK pipeline. The first two steps of the UNEAK pipeline (production of tag counts and merged tag counts) were run to produce a common starting point for both pipelines, requiring approximately 6h hours to run on the test environment from the raw sequence files. The UNEAK pipeline is not multi-threaded so the presence of 24 processors on this machine was not relevant. The remaining steps in the UNEAK pipeline took only 5min. Data from both UNEAK and Haplotag were filtered and formatted using the small helper-program “CbyT” described by Huang et al. (2014), which is now updated and provided in the current Haplotag distribution. The use of CbyT allowed parameters in either pipeline to be relaxed, such that data filtering could be tested at different levels from the same output. The total count of SNP loci from the UNEAK pipeline passing the population filter at a genotype-completeness threshold of >=50% was 12,780. At a threshold of >=80%, the count of filtered SNP loci was 4,260 (Table 1).

**Table 1.**
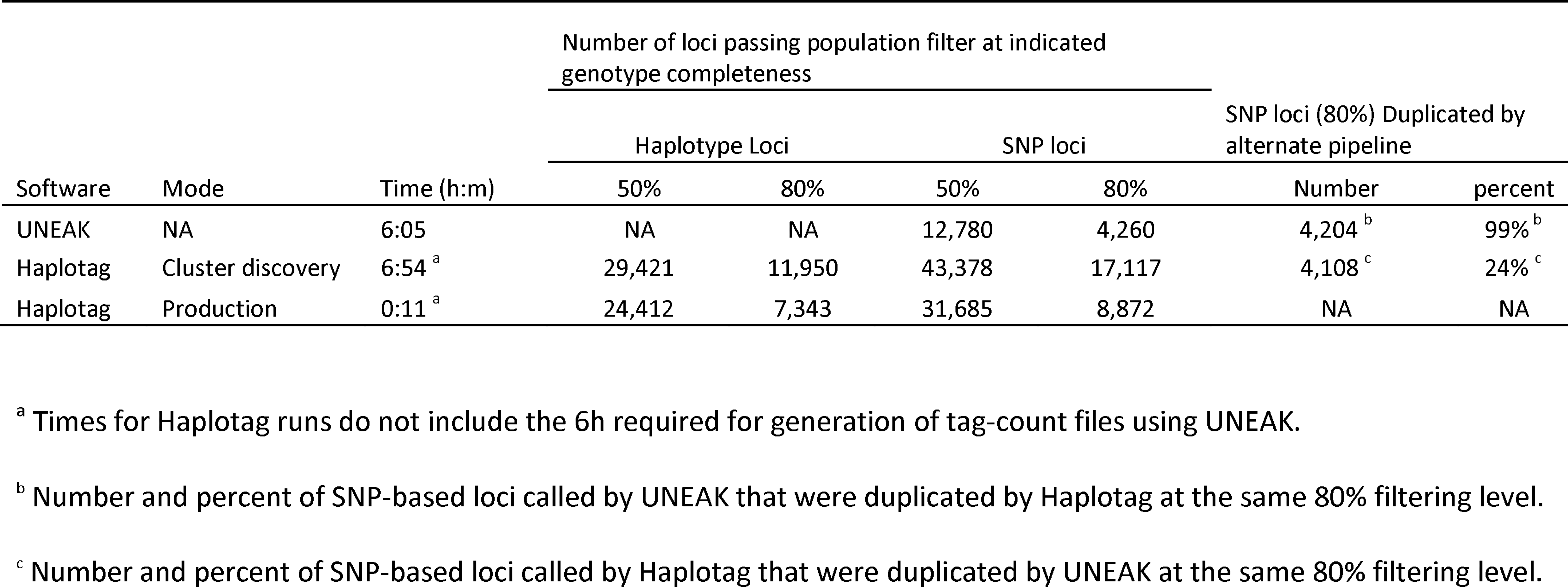
Comparison of GBS data analysis using UNEAK vs Haplotag software.

Running on the same machine, but utilizing 23 processors, the full Haplotag pipeline in cluster discovery mode took 6.9h in addition to the 6h required by UNEAK. The cluster discovery step took most of this execution time. After applying the same population filter, the number of Haplotag loci was 29,421 with a genotype-completeness of >=50% or 11,950 with a genotype-completeness of >=80%. When translated to SNP loci, the number of calls was 43,378 at >=50% completeness or 17,117 at >=80% completeness. The larger number of SNP loci relative to Haplotag loci is due to the presence of multiple SNP loci within some Haplotag loci.

In comparing the filtered SNP loci called by UNEAK to the SNP loci called from Haplotag, 4204 (99%) of the 4260 UNEAK SNPs filtered at >=80% genotype-completeness were identical to those called by Haplotag at the same filtering level. In contrast, UNEAK identified only 24% of the 17,117 Haplotag SNPs filtered at >=80% genotype-completeness. In general, Haplotag called most of the same SNP loci discovered by UNEAK, because these represented the clusters in Haplotag with exactly two haplotypes having only a single SNP difference. The small number of UNEAK SNPs that were missed by Haplotag are a result of rare haplotypes and/or sequencing errors that were aligned into a large cluster by Haplotag. In rare cases, this resulted in a complex cluster that was excluded from the Haplotag project because it exceeded the threshold for the maximum number of tags per cluster. The UNEAK pipeline has a different network-based strategy that is intended to exclude rare haplotypes, because it is designed to seek models with only two haplotypes and a single SNP. While it is possible to adjust Haplotag parameters to increase the coverage of UNEAK SNPs, this would be at the expense of a greater number of multi-haplotype models that are called by Haplotag.

Haplotag was also tested in production mode, which required only 11 minutes in our test environment. As shown in Figure 1, production mode uses loci and haplotypes discovered in a previous analysis to reduce the computation time and preserve an established nomenclature. When we used input files from the previously reported cluster-discovery run, we achieved exactly the same results, as expected. Thus, to test a different scenario, we used input files from an alternate analysis (not reported) where Haplotag had been run in haplotype discovery mode. In that analysis, clusters were built from the full set of SNP reference sequences reported by Huang et al. (2014), as well as from additional SNP reference sequences from subsequent work, encompassing a total of 3327 taxa. We had used this strategy in order to preserve SNP nomenclature with that of prior published and submitted work. Here, we wanted to test whether the haplotypes discovered using this large inventory of reference sequences would provide similar results to those achieved above using Haplotag in cluster discovery mode. The results of this analysis provided genotypes for 24,412 or 7,343 Haplotag loci (at >=50% or >=80% genotype-completeness, respectively), which translated to 31,685 and 8,872 SNP loci, respectively (Table 1). Averaged across filtering levels, this was a 33% reduction in called loci relative to those from Haplotag in full cluster discovery mode. The disadvantage of this strategy, which we have now demonstrated, is that the current production files have not incorporated a large number of high quality “new” SNPs that are discoverable only by Haplotag. This new result will be considered in future GBS work in oat, and will require careful addition of new clusters, loci, and haplotypes to the existing production files, while still preserving the legacy nomenclature.

Each Haplotag run produces a complete index of passport files, linking each Haplotag locus to a passport file for the cluster where that locus was called. While this index is written in HTML format, it can easily be manipulated into a table, which we have demonstrated in Supplement S3. This table provides links to the passport files for the 7,343 Haplotag loci called in the production mode and filtered at >=80% genotype-completeness. We have chosen this output because it contains legacy SNPs and nomenclature (from Huang et al. 2014) to which we have added known map positions. By loading all passport files to a web server, they do not need to be downloaded and duplicated by users of this resource. This strategy will be used in future to provide passports and metadata for public GBS data sets loaded into the T3/Oat database (https://triticeaetoolbox.org/oat/). Since passport files can also be saved and opened without the need for a web server, an individual passport file can easily be shared with a collaborator when there is an interest in inspecting the sequence and genotypes of a specific Haplotag locus.

### Limitations and future development

Haplotag was developed primarily to solve problems of genotyping in self-pollinating allopolyploid species without a reference genome. It will also function well in a self-pollinating diploid species. When paralogous loci exist, such that they are aligned together within the same cluster, Haplotag depends on a simple heterozygosity filter to build models of Haplotag loci that exclude haplotypes from non-homologous loci. Typically, this is very effective in self pollinating populations where heterozygotes are rare and this filter can be set at a low level (typically between 0.05 and 0.12). In populations where high rates of heterozygosity are expected (in F_2_ populations, or in populations of outcrossing species) a heterozygosity filter that was set higher (e.g. 0.65) could still be effective in excluding non-segregating haplotypes from paralogous locus, but complications could arise if multiple paralogous loci are segregating simultaneously. We initially considered the application of a Fishers’ test of contingency tables, but extending this test to an arbitrary number of haplotypes was beyond our programing skills. In future, we may consider adding additional population filters to expand the genetic scenarios in which Haplotag can be used, and we welcome suggestions in this regard.

## Supplementary Material

File S1. Complete user manual for Haplotag. Future updates may be available at http://haplotag.aowc.ca where the latest version of Haplotag software can also be downloaded.

File S2: Details of model selection from the passport file presented in Figure 2. A total of 57 models were evaluated, which represent all possible combinations with 2 or more members of the 6 potential haplotypes. Of these, 5 models met the filtering criteria. Model 42 was selected as the first valid locus with the greatest number of complete genotypes. Other models containing overlapping haplotypes from model 47 were then eliminated, and the process was iterated to select model 48 as a second valid locus.

Table S3. Index of haplotype based locus calls from the software Haplotag. Calls were made from primary sequence data originating from 894 taxa, described by Huang et al. (2014). Data were analysed in the Haplotag production mode, such that SNP nomenclature from the previous work was preserved.

